# PI3Kgamma promotes neutrophil extracellular trap formation by noncanonical pyroptosis in abdominal aortic aneurysm

**DOI:** 10.1101/2024.01.26.577503

**Authors:** Yacheng Xiong, Shuai Liu, Yu Liu, Jiani Zhao, Jinjian Sun, Baihong Pan, Wei Wang

## Abstract

**Background:** Abdominal aortic aneurysm (AAA) is one of the most life-threatening cardiovascular diseases while currently lacks effective drug treatment. NETs formation has been proved to be crucial trigger of AAA, so finding its upstream regulatory targets is the key to discovering therapeutic agents for AAA.

**Methods and Results:** we reveal that PI3Kgmma (PI3Kγ) is an upstream signal that regulates NETs formation. Inhibition of PI3Kγ reduces the expression of NETs and reduces inflammation in the aortic wall, thereby significantly ameliorating AAA. However, the mechanism of NETs formation regulated by PI3Kγ has not been fully elucidated. Using isolated bone marrow neutrophils, we show that PI3Kγ deficiency inactivates the noncanonical pyroptosis pathway (Capase11/GSDMD) to inhibit NETs expression, and subsequently found that PI3Kγ regulation of noncanonical pyroptosis via anchoring PKA is dependent on cAMP/PKA signaling, but not on classical PI3K/AKT signaling.

**Conclusions:** Our research uncovers the role and mechanism of PI3Kγ in AAA development and provides insights into AAA therapy from the perspective of NETs formation.

## Introduction

Abdominal aortic aneurysm (AAA) is a balloon-like bulge formed in the abdominal segment of the aortic wall caused by various factors, and its rupture is a major cause of death in adults^1^. Prevalence of AAA is up to 0.92% in people aged 30-79 worldwide^2^. Progressive aneurysmal dilatation eventually progresses to aneurysmal rupture, with a mortality rate of more than 81%^3^. So far, there is no effective drug to prevent the formation or growth of aneurysms in these patients^4^. Meanwhile, the existing drugs used in clinical trials for the treatment of AAA have not yet shown good efficacy^5^. Therefore, it is in desperate need to explore the potential mechanisms of AAA and discover the novel effective therapies.

Inflammation plays a pivotal role in the pathological process of AAA^6^. A variety of immune cells such as neutrophils, monocytes/macrophages, eosinophils, and lymphocytes infiltrate in the arterial wall tissue, which is associated with the development of AAA^7^. Among inflammatory cells, neutrophils are recruited to the aortic wall at early stage of AAA formation^8^. A prospective cohort study showed a strong positive association between neutrophil count and AAA development, independent of other traditional risk factors such as smoking, obesity, and atherosclerosis^9^. Limiting neutrophil recruitment to the arterial wall significantly reduced the incidence of elastase-perfusion AAA^10^, and inhibition of neutrophils using anti-neutrophil antibodies can be also effective in limiting the progression of experimental AAA^8^. In addition, neutrophil-derived biomarkers may be of clinical value for the monitoring and prognosis of AAA and may be used to guide early therapeutic intervention^11^, suggesting that neutrophils may be a therapeutic target of AAA. Nevertheless, the application of neutrophil-neutralizing antibodies to reduce neutrophils in the treatment of AAA has a short-term effect and damages the host’s innate immune defense function^12^. Therefore, elucidating the mechanism of neutrophils in promoting AAA is of great importance for finding new therapeutic drug targets.

Neutrophils play an important role in immune defense through phagocytosis, degranulation and formation of neutrophil extracellular traps (NETs)^13^. NETs are composed of depolymerized chromatin, citrullinated histones3 (Cit H3), granule proteins and cytoplasmic proteins^14^. NETs formation contributes to pathogens clearance; however, uncontrol NETs formation accelerates disease deterioration in sterile inflammatory diseases, including atherosclerosis^15^. Studies have shown that circulating NETs are strongly correlated with the severity of AAA^16^, and that NETs aggregate AAA via initiation and promotion of inflammatory responses, smooth muscle phenotype switching, and extracellular matrix degradation through matrix metalloproteinases (MMPs)^17–19^. Further study has proved that reducing NETs formation on abdominal aorta can markedly protect elastase-induced AAA^20^. Currently, blockage of NETs formation is achieved through inhibit peptidyl arginine deiminases (PADs) which are expressed in neutrophil and drive histone citrullination^21^. However, PAD-deficient mice exhibit innate immune dysfunction and are more susceptible to bacterial infections^22^. Therefore, identification of another possible trigger of NETs in AAA is critical for development of therapeutic agents for AAA^23^.

PI3Kγ belongs to the PI3K protein family, which is mainly expressed in immune cells and involved in the progression of inflammatory-related diseases. Blocking PI3Kγ is regarded as an effective strategy for the treatment of inflammatory diseases^24^. Thiago et al. first reported that the release of NETs from human neutrophils was dependent on the activation of PI3Kγ during Leishmania infection^25^. A recent study revealed that PI3Kγ inhibition could reduce NETs formation in the treatment of microscopic polyarteritis^26^, suggesting that PI3Kγ may be an upstream regulator of NETs formation. However, it is remains unclear whether PI3Kγ is involved in AAA progression by promoting NETs formation; moreover, the underlying mechanism of PI3Kγ regulating NETs is still poorly understood.

Herein, we revealed that PI3Kγ inhibition reduced neutrophil infiltration and NETs formation in the arterial wall of AAA. Reduction of NETs formation through PI3Kγ blockade is achieved through restraining noncanonical pyroptosis pathway. Regulatory role of PI3Kγ on noncanonical pyroptosis pathway accomplished through cAMP/PKA signaling pathway. The molecular mechanism of PI3Kγ regulating the progression of AAA, which is PI3Kγ-cAMP/PKA-noncanonical pyroptosis-NETosis, was clearly elucidated. Our study provides a new insight into the prevention of AAA and broadens the understanding of the upstream regulatory sites of NETs formation.

## Methods

### 2.1. reagents

The key reagents used in this experiment can be obtained in table S1

### 2.2. Ethics statement

This study was approved by the Ethics Committee of Central South University Xiangya Hospital (Approval No.201803481) in accordance with the Declaration of Helsinki principles. All animal procedures adhered to the US National Institutes of Health’s Guide for the Care and Use of Laboratory Animals.

### 2.3. Study design, subject recruitment, and sample collection

In the present study, AAA patients were recruited in Xiangya Hospital, Central South University (Changsha, China) from September 2021 to October 2022. AAA were comfirmed by computed tomography angiography (CTA) according to the 2018 American Society for Vascular Surgery (ASVS) guidelines for the diagnosis and treatment of AAA^27^. The AAA samples were collected from the patient receiving open aneurysm repair. The control tissue sample was derived from adjacent relatively normal abdominal aorta. The demographic and clinical variables can be obtained in table S2. All included subjects provided written informed consent.

### 2.4. Animal study

Male wildtype specific pathogen-free (SPF) C57BL/6J mice aged 8 weeks were purchased from SPF Hunan SJA Laboratory Animal Co., Ltd. (Changsha, China). PI3Kγ knockout mice (PI3K γ-/-) were purchased from GemPharmatech Co. Ltd. (Nanjing, China). Construction strategy and genotype identification are shown in Figure S1A-1B.

The Porcine pancreatic elastase (PPE) induced AAA model was established by intravascular PPE perfusion as previously described^28^. Briefly, PPE (4.0U/mL dissolved in sterile saline) was delivered into infrarenal aorta for 5min at 150mmHg pressure after mice were anesthetized with 1.5% isoflurane. The control group underwent sham surgery and were administrated with an identical volume of saline.

Cl-amidine (20 mg/kg) and H89 (2 mg/kg) was administered by intraperitoneal injection in PPE-induced AAA model at different endpoints.

Neutrophil adoptive transfer mice were established by tail vein injection of Neutrophils with 10^6^/mice once per day on days 2-3 after PPE models as described previously^20^. All AAA model mice were sacrificed for measurement of maximal aortic diameter and tissue collection on day 14 after the surgery. All animals were housed in SPF conditions at 23±1℃ with 50-60% humidity in a 12-hour light/dark cycle and were provided standard chow diet and water ad libitum.

### 2.5. Aortic ultrasound imaging

Abdominal aortas were visualized in isoflurane-anesthetized mice at study endpoints using the Color Doppler ultrasound system with ZS3 Exp Domain scanning imaging technique (Mindray). Cross-sectional images of the abdominal aortas were captured and maximal lumen diameters of the abdominal aortas at aneurysm sites were measured. All recordings were made by 2 researchers in a blinded manner.

### 2.6. Neutrophil isolation and culture

Mice were sacrificed and bone marrow was removed. Neutrophils from mice bone marrow were isolated using Mouse marrow neutrophil isolation kit (Solarbio, P8550). Immediately after isolation, the neutrophils layer was collected and resuspended in RPMI 1640 medium containing 5% fetal bovine serum and 1% penicillin/streptomycin.

The cells were incubated at 37℃ and 5% CO2 in a humidified incubator and used for NETosis studies after 30 minutes of isolation. The concentration of neutrophils was identified with staining and Flow cytometry in Figure S3A-3C.

### 2.7. Induction of neutrophil extracellular traps (NETs) and administration intervention in vitro

Freshly purified neutrophils were grown on Poly-L-Lysine-coated coverslips in a 24-well plate. For NETosis induction, different concentrations of LPS were used and the optimal concentration was 5 μg/mL (see Figure S3D-3E). Neutrophils were treated with PBS as a negative control and LPS (5 μg/mL, Sigma–Aldrich) or TNFa (50 ng/mL, MedChemExpress) for 4 h.

For the administration intervention of NETosis, neutrophils were pretreated with IPI549 (5 μM, MedChemExpress) or Disulfiram (30 μM, MedChemExpress) for 1 hour and then stimulated with LPS (5 μg/mL, Sigma–Aldrich) for 4 h to induce NET formation. In addition, to interfere with ROS or AKT pathways, neutrophils were pretreated with Diphenyleneiodonium chloride (DPI, 10 μM, MedChemExpress) or SC79 (4 μg/mL, MedChemExpress) for 1 hour after IPI549 administration, and then stimulated with LPS for 4 h to induce NET formation.

To explore pyroptosis signal during NETosis, neutrophils were primed with ultrapure LPS (100 ng/ml) for 3 hours, and subsequently stimulated with Nigericin (10 μM) for 1 hour after IPI549 administration to induce canonical inflammasome activation. For noncanonical inflammasome activation, neutrophils were primed with Pam3CSK4 (1 μg/ml) for 3 hours before IPI549 administration, and subsequently transfected without (Mock) or with ultrapure LPS (10 μg/ml) into the cytosol with DOTAP Liposomal Transfection Reagent (Roche) for 4 hours.

For the administration of cAMP/PKA inhibitor, H89 (20 μM, MedChemExpress) or MDL12330A (10 μM, MedChemExpress) were added to neutrophil for 1 hour after IPI549 administration, and then stimulated with LPS for 4 h to induce NET formation. After stimulation, cells were collected for subsequence test.

### 2.8. Immunohistochemistry

The aorta tissues were fixed in 4% paraformaldehyde, dehydrated in 70%-100% ethanol, embedded in paraffin and sectioned (5 μm). Antigen retrieval was performed using Sodium citrate-EDTA buffer in an autoclave for 6 min, followed by blocking nonspecific binding with 3% BSA containing 0.3% Triton X-100 for 1 hour. Tissue sections were incubated with primary antibodies (see Table S1) overnight at 4℃. Subsequently, the sections were rinsed and incubated with HRP-conjugated secondary antibodies according to the manufacturer’s protocol, then staining with DAB (ZSGB-BIO, ZLI-9018) and hematoxylin. Images were obtained by a Leica DM6 M.

### 2.9. Immunofluorescence staining

The steps of tissue immunofluorescence before primary antibody incubation were identical to those of tissue immunohistochemistry. Cells for immunofluorescence were fixed with 4% paraformaldehyde for 15 min, washed with PBS and permeabilized with 0.3% Triton X-100 for 10 min. After blocking with 3% BSA for 1 hour. Tissue sections or cells were incubated with primary antibodies overnight at 4℃ and then incubated with corresponding fluorescent-conjugated secondary antibodies for 1 hour at room temperature. The antibodies used are listed in Table S1. Images were obtained by a Leica DMi8 M (Leica Microsystems, Germany). Cells were stained with DAPI, citrullinated histone 3 (Cit H3) antibody and myeloperoxidase (MPO) antibody to detect NETs.

### 2.10. VVG staining

The aorta tissues were harvested and fixed with 4% PFA before paraffin embedding. Tissue sections (5μm) were processed for Verheff’s Elastic staining using Verhoeff Van Gieson (VVG) Elastic Stain Kit (Abiowell, AWI0267b) according to the manufacturer’s protocol. Images were obtained by a Leica DM6 M. The severity of elastin fragmentation was evaluated and graded as follows: grade 1, no degradation; grade 2, mild elastin degradation; grade 3, moderate elastin degradation; grade 4, severe elastin degradation. Each sample was assessed by 2 independent pathologists.

### 2.11. Wright-Giemsa staining

Purified neutrophils were smeared and stained in accordance with the manufacturer’s instruction of Wright-Giemsa stain solution (Solarbio, G1020), and images were obtained by a Leica DM6 M.

### 2.12. Flow cytometry

Purified neutrophils were resuspended in cell staining buffer and distribute per 100μL of cell suspension (1 x 10^6^cells/tube), then incubated with Zombie Aqua^TM^ dye (BioLegend, 423101) for 20 mins in the dark at room temperature. Cells was washed with PBS and preincubated with 5 ml of TruStain FcX™ PLUS (anti-mouse CD16/32) Antibody (Fc Receptor Blocking Solution, BioLegend, 156603) for 10 minutes on ice to reduce nonspecific immunofluorescent staining. Then, the cell suspension was incubated with FITC-conjugated fluorescent antibody against CD11b (BioLegend, 101205) and APC-conjugated fluorescent antibody against Ly6G (BioLegend, 127613) on ice for 20 minutes in the dark (see Table S1). After washing with PBS twice, flow cytometry was conducted using a Cytek DxP Athena^TM^ flow cytometer (Cytek Biosciences) and the data were analyzed using FlowJo 10 software.

### 2.13. cAMP concentration

The cAMP concentration of tissues and neutrophils were measured by Mouse cAMP ELISA Kit (Jianglai biotech, JL13362) in accordance with the manufacturer’s instruction.

### 2.14. PKA Kinase Activity

Purified neutrophils and tissue PKA activities were measured by the nonradioactive PKA Kinase Activity Assay Kit (Abcam, ab139435) following manufacturers’ instructions.

### 2.15. Quantitative RT-PCR

Total RNA was extracted using RNAiso Plus reagent (Takara, 9019) and reverse transcribed into cDNA using the Hifair® AdvanceFast One-step RT-gDNA Digestion SuperMix (Yeasen, 11151ES60). DNA was amplified under the following reaction conditions: denaturing at 95℃ for 3 minutes, followed by 40 cycles of annealing at 95℃ for 5 s and extension at 60℃ for 31 s in the presence of 2X Universal SYBR Green Fast qPCR Mix (Abclonal, RK21203) and primers. The PCR products were detected using an Applied Biosystems ViiA 7 Real-Time PCR System (ThermoFisher Scientific). Relative gene expression was determined by the 2^-ΔΔCt^ method. Primers were designed according to the coding strand of genes in the National Center for Biotechnology Information (NCBI) database and are listed in Table S3.

### 2.16. Western blot

For cleaved IL-1β of cell supernatant, the proteins purified was conducted by Methanol-Chloroforms Extraction method as previous described^29^. Briefly, neutrophil suspensions were centrifuged at 2000rpm for 5 mins and cell supernatants were collected. 500 μL methanol and 125μL chloroform were added into 500μL of cell supernatants, and then vigorously mixed. After centrifugation (13000 rpm, 5mins), the upper clear liquid was removed, leaving the white pellet untouched. At this time, 500 μL of methanol was added and the white pellet was blown and dispersed into small pieces. With another round of centrifugation (13000 rpm, 5 min), the white pellet was left for drying at 55℃ for 10 mins until no visible liquid was evident. Finally, 1xSDS loading buffer was added to the dry proteins and heated for 5 min at 95°C, followed by immunoblotting analyses. For cell or tissue samples, proteins were isolated by RIPA buffer (Beyotime, P0013B) containing Protease and phosphatase inhibitor (Beyotime, P1045) on ice, and their concentrations were determined using a BCA protein assay kit (Beyotime, P0012). After adding SDS-PAGE protein loading buffer (Beyotime, P0015L) and denaturing at 95℃ for 10 minutes, equal amounts of protein samples were separated in a SDS-PAGE gel and transferred to PVDF membranes. The membrane was blocked with 5% nonfat milk in TBST buffer for 1 h at room temperature, incubated with appropriate primary antibodies overnight at 4℃ and then with horseradish peroxidase-conjugated secondary antibody for 1h at room temperature. The protein bands were visualized using an Immobilon Western HRP substrate Luminol Reagent (Millipore, WBKLS0500) and ChemiDoc XRS+ Imaging system (Bio-Rad). Densitometry analysis was performed using Image J software. The antibodies used in the present study are listed in Table S1.

### 2.17. Statistical analysis

Data statistical analysis was performed using Prism 9.5 software (GraphPad Software, Inc.). Values were shown as mean ± standard deviation (SD). The student’s t test was utilized between two groups and one-way ANOVA was used between more than two group comparison. Multiple groups with two variables were evaluated by two-way ANOVA followed by Bonferroni test. For different intervention between multiple groups, statistical analysis was performed using one-way ANOVA followed by Fisher’s least significant difference (LSD) post hoc test. For all statistical methods, P < 0.05 were considered significance: *P < 0.05, **P < 0.01, ***P <0.001.

## Results

### Neutrophil infiltration and NETs formation are upregulated in both human and mouse AAA tissue

To reveal levels in neutrophil infiltration and NETs formation in AAA wall, we measured the neutrophil numbers and quantified NETs both in human AAA samples and control adjacent abdominal aortic tissue (adjacent AA). The results showed that the degree of neutrophil infiltration in AAA was significantly more severe compared with those in control subjects (Figure 1A and 1B). Moreover, immunofluorescence staining of the aortic tissue revealed that increased NETs expression was mostly observed in AAA in comparison with that of control (Figure 1C and 1D). Subsequence, this image results was further verified at the protein level. The expression of Cit H3 protein, a marker of NETs, was markedly upregulated in AAA (Figure 1E and 1F). To further confirmed the above results, we repeated the experimental in mouse model of PPE-induced AAA. A similar increased of neutrophil infiltration and NETs formation was observed in AAA compared with aortas from saline-treat mice (Figure 1G-1J). To quantify the timing of NETs formation, protein from abdominal aortas were collected at 0, 3, 7 and 14 days after elastase perfusion. Cit H3 expression peaked in aortas during day 3 to 7 and downregulated slowly at day 14 (Figure 1K and L). These results all suggest that the abnormal neutrophil infiltration and NETs expression are closely associated with AAA.

**Figure 1.**
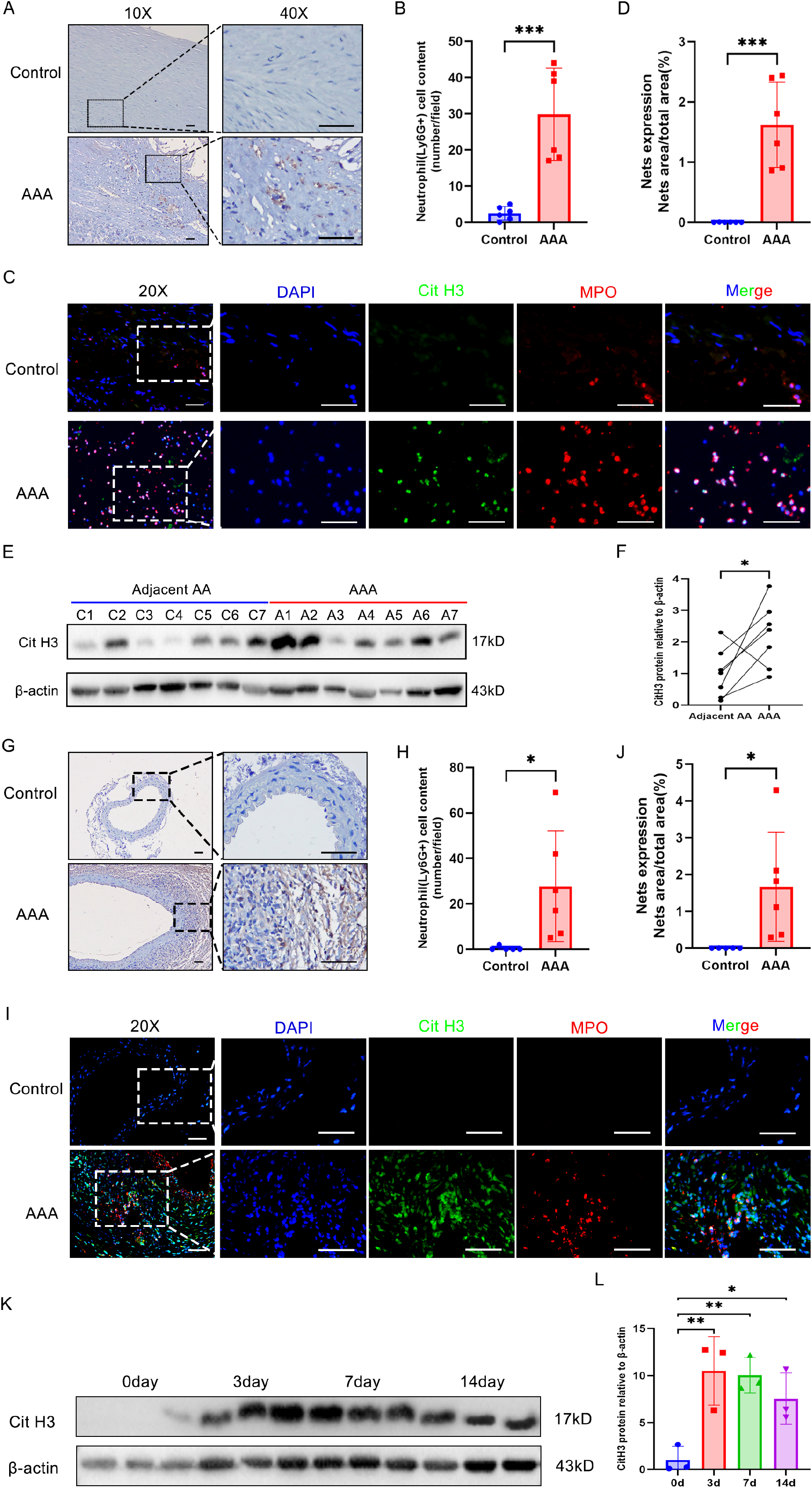
Neutrophil and NETs are upregulated in both human and mouse AAA tissue (A and B) Representative immunohistochemistry staining images (A) and quantitative comparison (B) of neutrophil infiltration in human abdominal aortic sections. Neutrophil infiltration was assessed by Ly6G+ neutrophil numbers per high-power field (40X). n=6. Scale bars, 50μm. (C and D) Representative immunofluorescence staining images (C) and quantitative comparison (D) of NETs in human abdominal aortic sections. Nets expression was calculated by NET area/total area (%) according to the fluorescence colocalization of DNA (DAPI, Blue), Citrullinated histone 3 (Cit H3, Green), and Myeloperoxidase (MPO, red). n=6. Scale bars, 50μm. (E and F) Representative western blot image (E) and quantitative analysis (F) of Cit H3 protein expression in human AAA tissue and adjacent relatively normal abdominal aortic tissue. n = 7. (G and H) Representative immunohistochemistry staining images (G) and quantitative comparison (H) of neutrophil infiltration in mouse abdominal aorta after PPE-induced AAA. Neutrophil infiltration was assessed by Ly6G+ neutrophil numbers per high-power field (40X). n=5 in control group and n=6 in AAA group. Scale bars, 50μm. (I and J) Representative immunofluorescence staining images (I) and quantitative comparison (J) of NETs in mouse abdominal aorta after PPE-induced AAA. Nets expression was calculated by NET area/total area (%) according to the fluorescence colocalization of DNA (DAPI, Blue), Citrullinated histone 3 (Cit H3, Green), and Myeloperoxidase (MPO, red). n=5 in control group and n=6 in AAA group. Scale bars, 50μm. (K and L) Representative western blot image (K) and quantitative analysis (L) of Cit H3 protein expression in mouse AAA tissue at different time points after PPE surgery. n = 3. (B, D, H, and J) Two-tailed unpaired student’s t test. (F) Two-tailed paired student’s t test. (L) One-way ANOVA followed by Fisher’s least significant difference (LSD) post hoc test.

### Inhibition of NETs alleviates elastase-induced AAA in mice

To reveal the crucial effect of NETs formation in AAA development. We delivered Cl amidine (A PAD4 inhibitor, blocking NETs formation) on the cure effect on PPE-induced AAA (Figure 2A). Analysis of ultrasonography of the abdominal aorta demonstrated smaller aortas lumen diameter in the Cl amidine administration group than in the vehicle (saline) group from day 7 and 14 after PPE surgery (Figure 2B and 2C). There were also significant differences in the external diameter of aorta between the Cl amidine and vehicle groups (Figure 2D and 2E). Histomorphology analysis showed that the degradation of elastin in the Cl amidine group was minder than that in the control group (Figure 2F and G). In order to verify the inhibitory effect of Cl amidine, immunofluorescence staining of NETs formation in aorta were conducted and results showed that NETs expression was decreased markedly after Cl amidine administration treatment (Figure 2H and I). In addition, the Cit H3 protein was significantly reduced at any time point after PPE surgery in Cl amidine mice (Figure 2J and K). Previous studies demonstrated that NETs exacerbated AAA by promoting smooth muscle phenotypic switching, inflammatory release, and matrix degradation^18^. We also observed the similar results. The mRNA levels of VSMC contractile gene Tagln, Cnn1, Acta2 and Myh11 were increased while synthetic gene Klf4 and Spp1 was reduced in Cl amidine group. Additionally, the proinflammatory genes (IL6, TNFa, Ccl2, IL-1 β) and the matrix metalloproteinase genes MMP2 and MMP9, were significantly decreased in the abdominal aorta of Cl amidine treated mice compared with those of control (Figure 2L). These data indicate that blocking the formation of NETs can effectively inhibit the progression of AAA.

**Figure 2.**
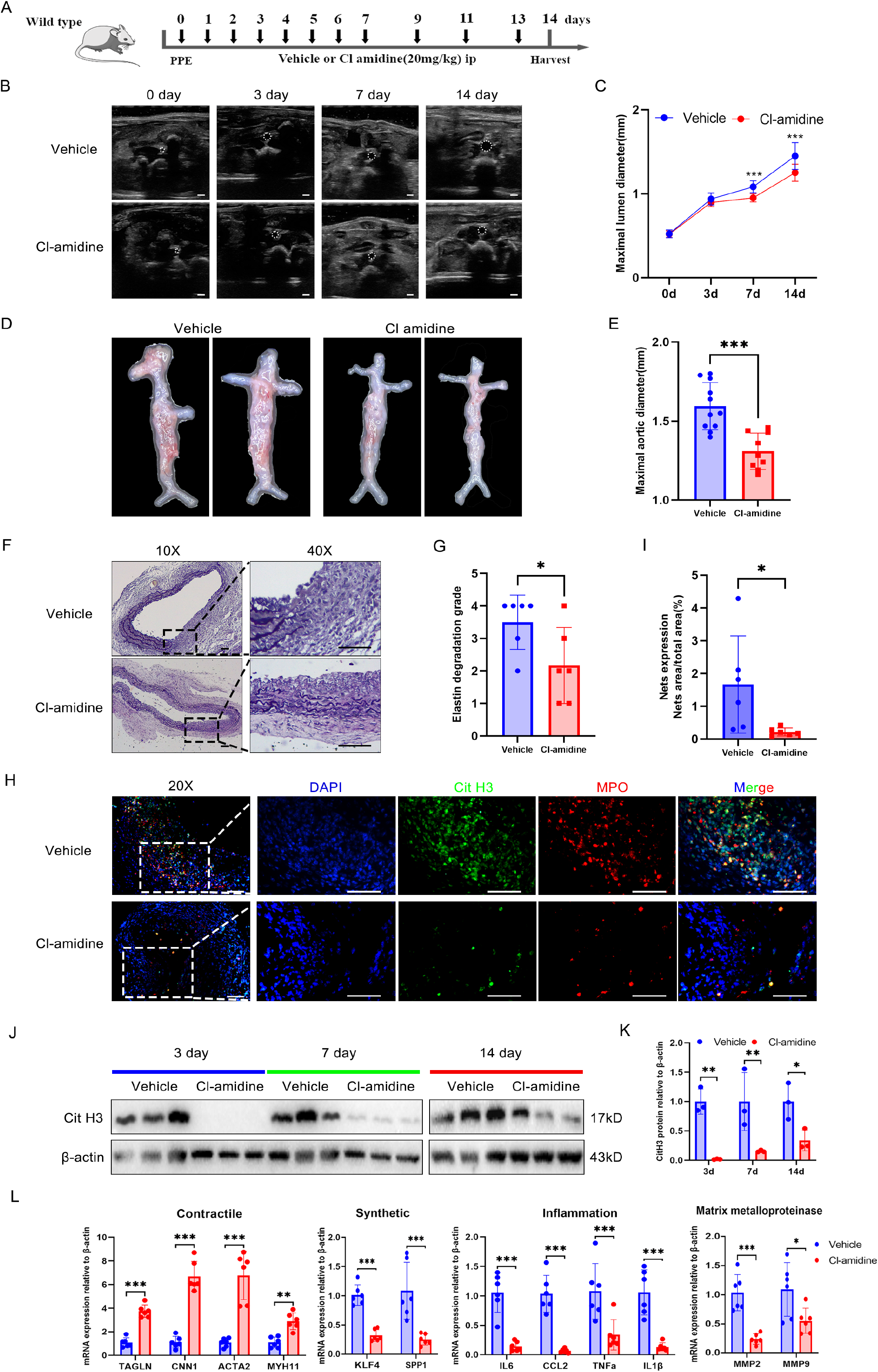
Inhibition of NETs alleviates elastase-induced AAA in mice (A) Schematic diagram of the animal study to verify the effect of cl amidine administration on PPE-induced AAA. (B and C) Representative ultrasonic images (B) and quantitative comparison (C) of maximal lumen diameter in mouse abdominal aorta at different time points with various treatments. n=13 in Vehicle group and n=15 in cl amidine group. Scale bar: 1 mm (D and E) Representative macroscopic images (D) and quantitative comparison (E) of maximal aortic diameter in mouse abdominal aorta at 14 days after PPE-induced AAA. n=11 in Vehicle group and n=9 in cl amidine group. (F and G) Representative VVG staining images (F) and quantitative comparison (G) of mouse abdominal aorta in PPE-induced AAA. n = 6. Scale bars, 50 μm. (H and I) Representative immunofluorescence staining images (H) and quantitative comparison (I) of NETs in mouse abdominal aorta after PPE-induced AAA. Nets expression was calculated by NET area/total area (%) according to the fluorescence colocalization of DNA (DAPI, Blue), Citrullinated histone 3 (Cit H3, Green), and Myeloperoxidase (MPO, red). n=6. Scale bars, 50μm. (J and K) Representative western blot image (K) and quantitative analysis (L) of Cit H3 protein expression in mouse AAA tissue at different time points with various treatment. n = 3. (L) Relative mRNA levels of contractile, synthetic, inflammation, and matrix metalloproteinase genes in mouse abdominal aorta after PPE-induced AAA. n=6. (C, K, and L) Two-way ANOVA followed by Bonferroni test. (E, G, and I) Two-tailed unpaired student’s t test. (F) Two-tailed paired student’s t test.

### PI3Kγ knockout reduces neutrophil infiltration and NETs release and improve AAA

Giving that inhibition of PAD impair innate immune defense system^22^, it is crucial to explore upstream regulator factors of NETs formation. PI3Kγ is involved in the regulation of neutrophil function^30^. We speculated that PI3Kγ participate in the development of AAA via inducing neutrophil NETs formation. Therefore, PI3Kγ knockout mice (PI3Kγ-/-) was generated (Figure S1A and S1B). Analysis on the PI3Kγ expression of abdominal aorta showed a significant increase in PI3Kγ protein of WT mice on day 14 after elastase perfusion compared to day 0 (Figure S1C). To further elucidate the impact of PI3Kγ on AAA, we performed PPE-induced AAA on PI3Kγ-/- mice and WT mice. Results revealed that PI3Kγ deficiency exhibited the similar effect on alleviating PPE-induced AAA as Cl amidine administration. Analysis of the ultrasonography, gross appearance, and elastic Van Gieson staining of the abdominal aorta demonstrated PI3Kγ-/- mice had milder aortas compared to WT mice (Figure 3A–3D and Figure S2A-S2B). These results suggested that PI3Kγ is an important etiological factor of AAA. Meanwhile, neutrophil infiltration and NETs formation were reduced in the PI3Kγ-/- group compared to the WT group (Figure 3E–3H). Furthermore, the expression of Cit H3 protein in PI3Kγ-/- mice was significantly lower than that in WT mice (Figure 3I–J). These results suggested that PI3Kγ is the upstream regulatory molecule of NETs formation. To determine the downstream signals of NETs as mentioned above, the real-time quantitative PCR was conducted. There was no significant difference in Cnn1, Acta2, and Klf4 of VSMC marker between the WT and PI3Kγ-/- groups; however, there were significant differences in Tagln, Myh11, and Spp1 between the two groups. The inflammatory gene, including IL6, TNFa, Ccl2, IL-1β were significantly downregulated in PI3Kγ-/- mice. In addition, it was MMP9 but not MMP2 that was reduced in PI3Kγ-/- mice (Figure S2C). All these results suggest that PI3Kγ blockade may alleviate the development of AAA by reducing the formation of NETs.

**Figure 3.**
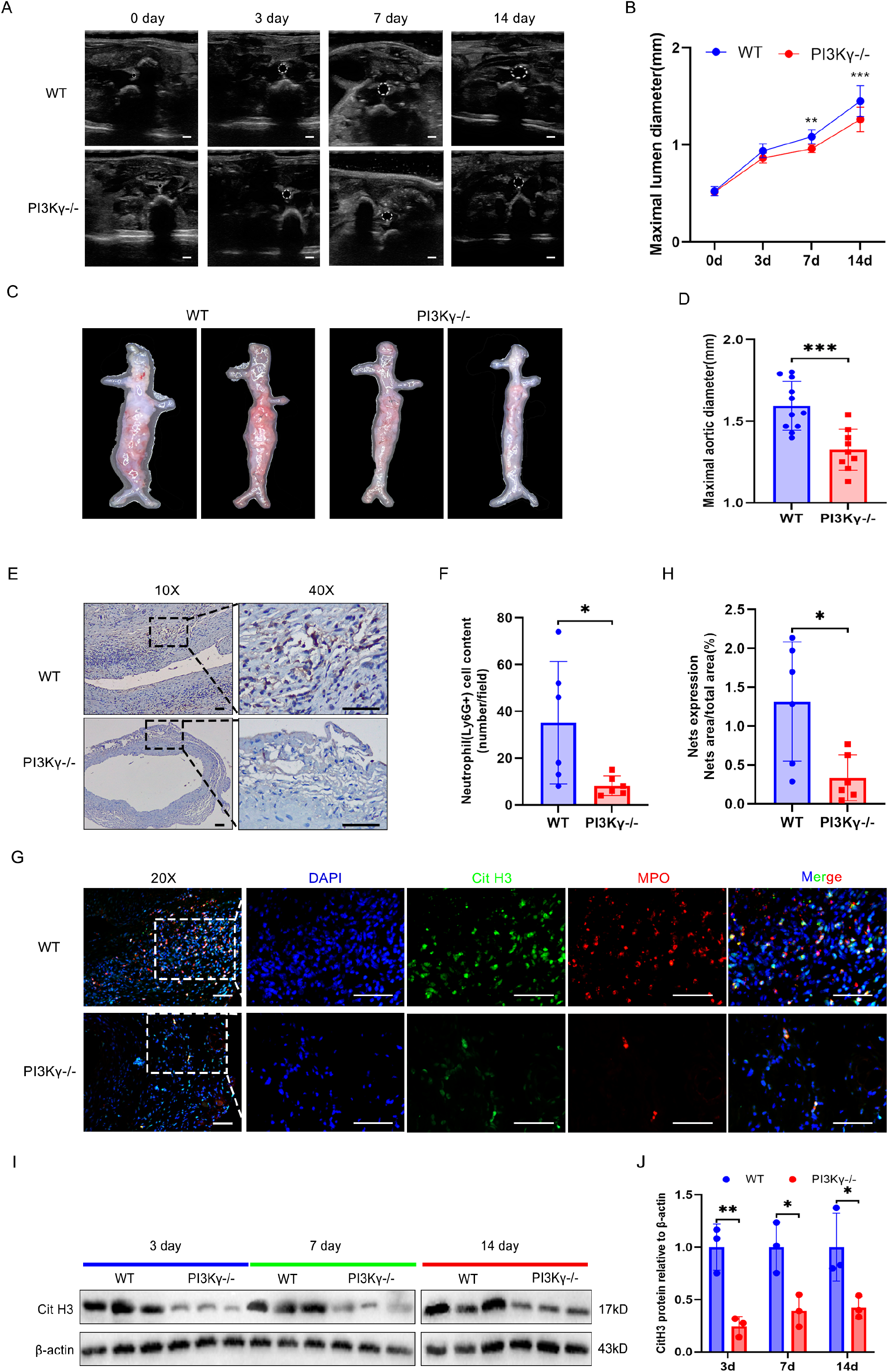
PI3Kγ knockout reduces neutrophil infiltration and NETs formation and improve AAA (A and B) Representative ultrasonic images (A) and quantitative comparison (B) of maximal lumen diameter in mouse abdominal aorta at different time points after PPE-induced AAA. n=13. Scale bar: 1 mm (C and D) Representative macroscopic images (C) and quantitative comparison (D) of maximal aortic diameter in mouse abdominal aorta at 14 days after PPE-induced AAA. n=11 in wild type (WT) group and n=9 in PI3Kγ-/- group. (E and F) Representative immunohistochemistry staining images (E) and quantitative comparison (F) of neutrophil infiltration in mouse abdominal aorta after PPE-induced AAA. Neutrophil infiltration was assessed by Ly6G+ neutrophil numbers per high-power field (40X). n=6. Scale bars, 50 μm. (G and H) Representative immunofluorescence staining images (G) and quantitative comparison (H) of NETs in mouse abdominal aorta after PPE-induced AAA. Nets expression was calculated by NET area/total area (%) according to the fluorescence colocalization of DNA (DAPI, Blue), Citrullinated histone 3 (Cit H3, Green), and Myeloperoxidase (MPO, red). n=6. Scale bars, 50 μm. (I and J) Representative western blot image (I) and quantitative analysis (J) of Cit H3 protein expression in mouse AAA tissue at different time points after PPE surgery. n = 3. (B, L, and M) Two-way ANOVA followed by Bonferroni test. (F, H, and J) Two-tailed unpaired student’s t test.

### Deficiency of PI3Kγ inhibits NETs formation in neutrophil

To clarify the causal relationship between PI3Kγ and NETs formation in neutrophils. Neutrophil was isolated and purified (purity > 90%) from mice (Figure S3A-S3C). Then, we stimulated the isolated neutrophils with LPS which has been confirmed as the classical inducer of NETs^31^. We tested the induction of NETs formation by different concentrations of LPS and found that the most appropriate concentration was 5 μg/mL (Figure S3D-S3E). We found that LPS-treated neutrophils from WT mice formed large amounts of NETs. Moreover, IPI549 (PI3Kγ inhibitor) could significantly reduce NETs formation induced by LPS (Figure S4A-S4D). In addition, to mimic an inflammatory state of AAA, isolated neutrophils were also stimulated by TNFa. Results revealed that NETs formation was significantly increased, but was reversed after IPI549 administration (Figure S4E-S4H). To further corroborate the experimental results, we repeated most of the above experimental in neutrophils purified from PI3Kγ-/- and WT mice, the results were completely consistent with those of PI3Kγ inhibitor treatment (Figure 4A-4H). All the results suggested that inhibition of PI3Kγ in neutrophils could reduce NETs formation.

**Figure 4.**
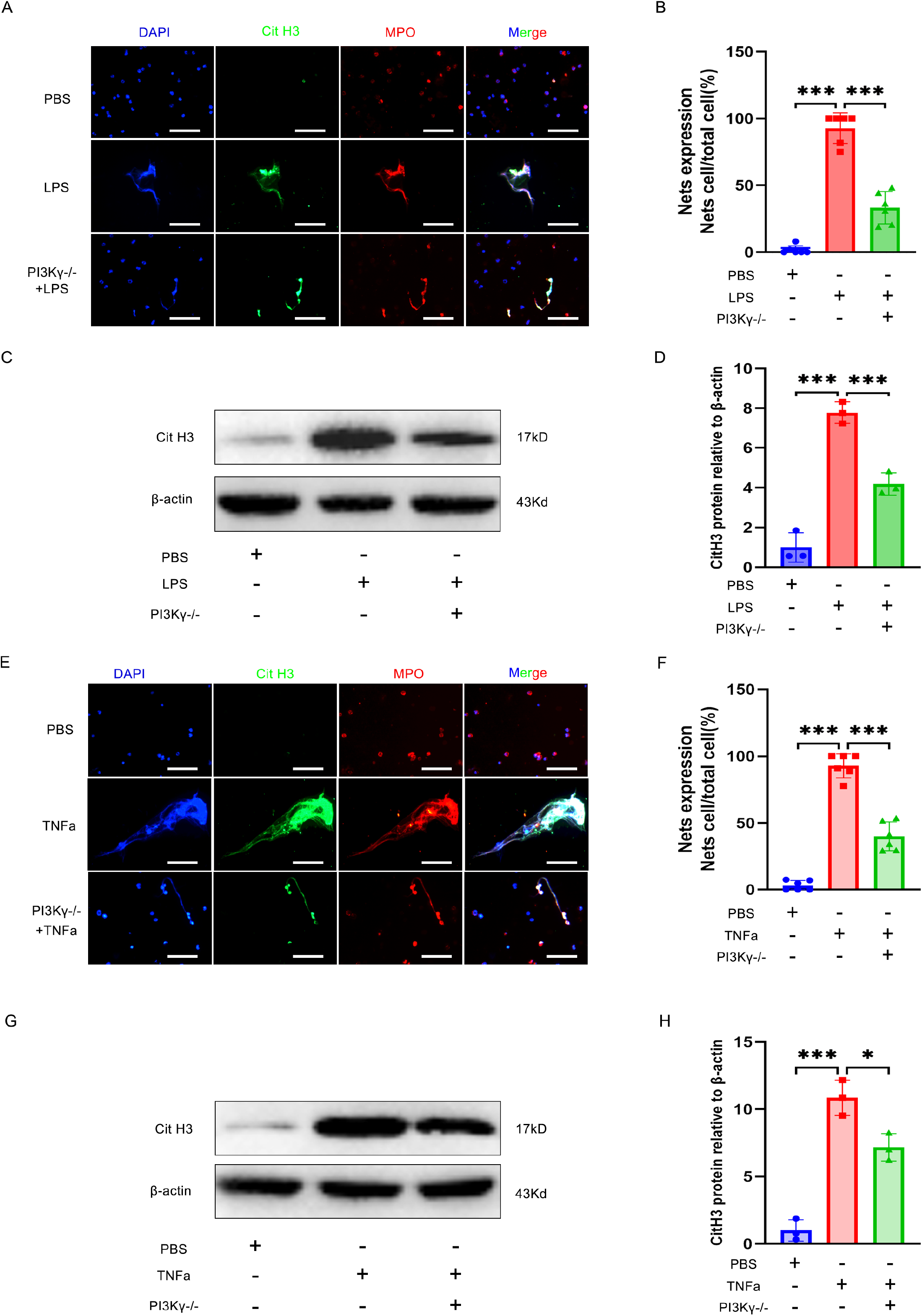
Deficiency of PI3Kγ inhibits NETs formation in neutrophil (A, B, E, and F) Representative immunofluorescence staining images (A, E) and quantitative comparison (B, F) of NETs produced by neutrophils with different treatments: PBS (WT-derived neutrophils), LPS (5 μg/mL, WT-derived neutrophils), LPS (PI3Kγ-/- derived neutrophil); PBS (WT-derived neutrophils), TNFa (50 ng/mL, WT-derived neutrophils), TNFa (PI3Kγ-/- derived neutrophil). NETs were detected using immunofluorescent staining of DNA (DAPI, blue), Citrullinated histone 3 (Cit H3, Green), and Myeloperoxidase (MPO, red). NETs expression was calculated by NETs cell numbers/total cell numbers per high-power field (40X). n=6. Scale bars, 50 μm. (C, D, G, and H) Representative western blot images (C, G) and quantitative comparison (D, H) of Cit H3 protein expression in each group of neutrophils with different treatments as described in (A), (B), and (G) or (C), (D), and (H). n=3. (B, D, F, and H) One-way ANOVA followed by Fisher’s LSD post hoc test.

### PI3Kγ expression in neutrophil is required for NETs formation and AAA progression in mice

To further reveal the role of PI3Kγ from neutrophils in AAA formation, we generated elastase-induced AAA in PI3Kγ-/- mice. Since neutrophil was the only source of PI3Kγ in PI3Kγ-/- mice, adoptive transfer of neutrophils isolated from WT mice into PI3Kγ-/- mice was performed. A Schematic diagram of the process of neutrophil adoptive transfer experiment is shown in Figure 5A. Results indicated that PI3Kγ-/- mice receiving WT neutrophils exhibited more severe AAA lesions compared with PI3Kγ-/- mice without neutrophil transfer. Both the maximal lumen and external diameter of the abdominal aorta of neutrophil-transferred PI3Kγ-/- mice were significantly greater than those of vehicle-treated PI3Kγ-/- mice (Figure 5B-5E). Verhoeff-Van Gieson staining of the abdominal aorta and found that the degree of elastin degradation was more severe in neutrophil-transferred PI3Kγ-/- mice (Figure 5F-5G). Subsequence, NETs formation was evaluated by immunofluorescence and Western blot, results indicated that NETs expression was significantly increased in neutrophil-transferred PI3Kγ-/- group than in vehicle treated PI3Kγ-/- group (Figure 5H-5K). Furthermore, adoptive transfer of neutrophil significantly decreased the expression of VSMC contractile genes, for example, Acta2, Cnn1, Tagln, and Myh11. Significantly increased expression of genes associated with synthetic and inflammatory phenotype and extracellular matrix degradation was also observed in neutrophil-transferred PI3Kγ-/- group compared with that in vehicle treated PI3Kγ-/- group (Figure 5L). These results suggest that neutrophil-derived PI3Kγ is the key regulatory molecular on NETs formation in mouse model of PPE-induced AAA.

**Figure 5.**
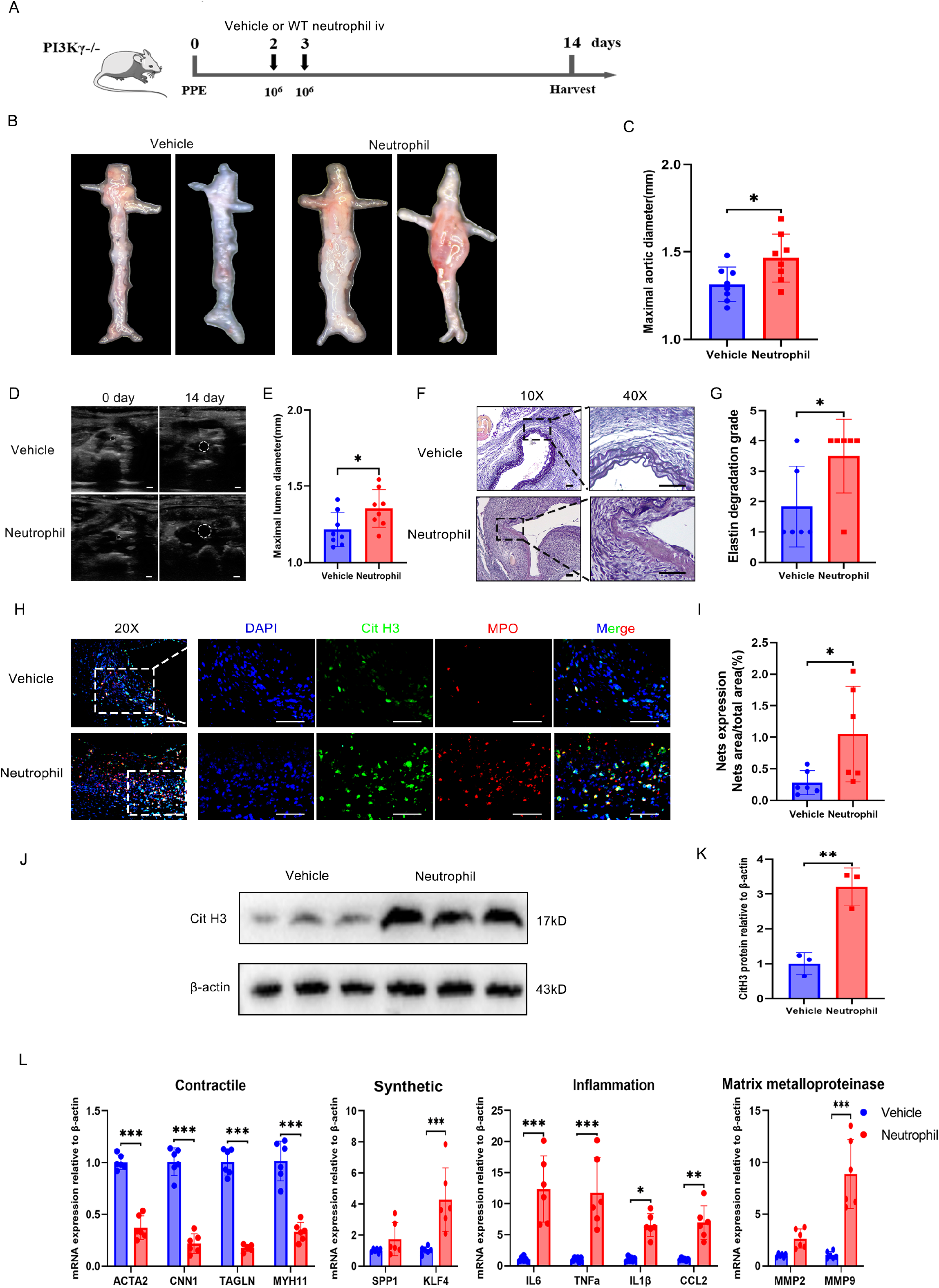
PI3Kγ expression in neutrophil is required for NETs formation and AAA progression in mice (A) Schematic diagram of the experimental process of neutrophil adoptive transfer in PPE-induced AAA PI3Kγ-/- mice. (B and C) Representative macroscopic images (B) and quantitative comparison (C) of maximal aortic diameter in PI3Kγ-/- mice after PPE surgery. n=8. (D and E) Representative ultrasonic images (D) and quantitative comparison (E) of maximal lumen diameter in PI3Kγ-/- mice after PPE surgery. n=8. Scale bar: 1 mm (F and G) Representative VVG staining images (F) and quantitative comparison (G) of VVG staining score in PI3Kγ-/- mice after PPE surgery. n = 6. Scale bars, 50 μm. (H and I) Representative immunofluorescence staining images (H) and quantitative comparison (I) of NETs in PI3Kγ-/- mice abdominal aorta after PPE-induced AAA. Nets expression was calculated by NET area/total area (%) according to the fluorescence colocalization of DNA (DAPI, Blue), Citrullinated histone 3 (Cit H3, Green), and Myeloperoxidase (MPO, red). n=6. Scale bars, 50 μm. (J and K) Representative western blot image (J) and quantitative analysis (K) of Cit H3 protein expression in the abdominal aorta of PI3Kγ-/- mice. n = 3. (L) Relative mRNA levels of contractile, synthetic, inflammation, and matrix metalloproteinase genes in the abdominal aorta of PI3Kγ-/- mice. n=6. (C, E, G, I and K) Two-tailed unpaired student’s t test. (L) Two-way ANOVA followed by Bonferroni test.

### PI3Kγ promotes NETs formation via noncanonical pyroptosis pathways in vitro

Our results described above suggested that inhibition of PI3Kγ alleviated AAA through suppressing NETosis. Therefore, the following study was designed to elucidate the molecular mechanism of PI3Kγ on regulating NET formation. Previous studies have identified reactive oxygen species (ROS) as critical for the induction of NETs formation^32^. Western blot was conducted to detect the protein expression of NADPH oxidase 2 (NOX2, a key enzyme in inducing ROS production). Results revealed that LPS-treated neutrophils released large amounts of NETs, whereas PI3Kγ inhibition or DPI administration (a specific NOX inhibitor) could significantly reduce NETs formation, and they also had an obvious synergistic effect. Interestingly, there was no difference in NOX2 protein expression during PI3Kγ blockage (Figures S5A–S5F). These results suggest that the inhibitory effect of PI3Kγ deficiency on NET formation in neutrophils is independent of ROS signaling.

Previous studies have suggested that pyroptosis is involved in the formation of NETs^33^, so we measured the expression levels of Gasdermin D (GSDMD, a key protein in pyroptosis signaling pathway). Results indicated that the levels of N-GSDMD protein, the active splicing form of GSDMD, was significantly increased in human AAA compared with adjacent abdominal aorta tissue (Figure S6A-S6B). Furthermore, WT mouse neutrophils were isolated and treated with LPS. The results showed that LPS induced significant increase in N-GSDMD and Cit H3 protein, which were reversed by disulfiram (a specific GSDMD inhibitor) administration (Figures 6A–6C). These results indicates that pyroptosis is a key upstream regulatory signal for NETs formation. To investigate whether PI3Kγ regulates NETs formation through pyroptosis pathway, we tested the expression of pyroptosis pathway-related proteins in WT or PI3Kγ-/- neutrophil. Results showed that the expression of IL-1 β and N-GSDMD was significantly inhibited after PI3K γ deficiency (Figures 6D–6F). These findings suggested that pyroptosis is a key bridge for PI3Kγ to promote NETs formation. Previous studies have shown that the occurrence of pyroptosis depends on the activation of canonical or noncanonical inflammatory pathway^34^. Subsequently, tests were carried out to examine whether canonical or noncanonical inflammatory pathways are involved in PI3Kγ-induced NETs. In order to activate canonical inflammatory pathway, neutrophils from WT or PI3Kγ-/- mice were primed with LPS followed by administration of nigericin to activate the NLRP3 inflammasomes (Caspase1). In order to activate noncanonical inflammatory pathway, Pam3CSK4 (a TLR1/2 agonist) was utilized to suppress basal apoptosis, then LPS was transfected into the neutrophil cytosol to activate caspase 11. The results indicated that both canonical and noncanonical inflammasome activation significantly increased CitH3 expression and IL-1β release. However, only the Caspase11/GSDMD signaling (noncanonical pyroptosis pathway) was inhibited after PI3Kγ blockade, whereas no significant difference in canonical pyroptosis pathway associated proteins (Caspase1/IL-1β) were observed between PI3Kγ-/- and WT groups (Figure 6G-6L). Subsequently, we repeated the above experiment in neutrophils isolated from WT mice by using PI3Kγ inhibitor (IPI549). The results were completely consistent with those from PI3Kγ-derived neutrophils (Figure S6C-S6K). Our data suggested that PI3Kγ might promote NETs formation via noncanonical pyroptosis pathways.

**Figure 6.**
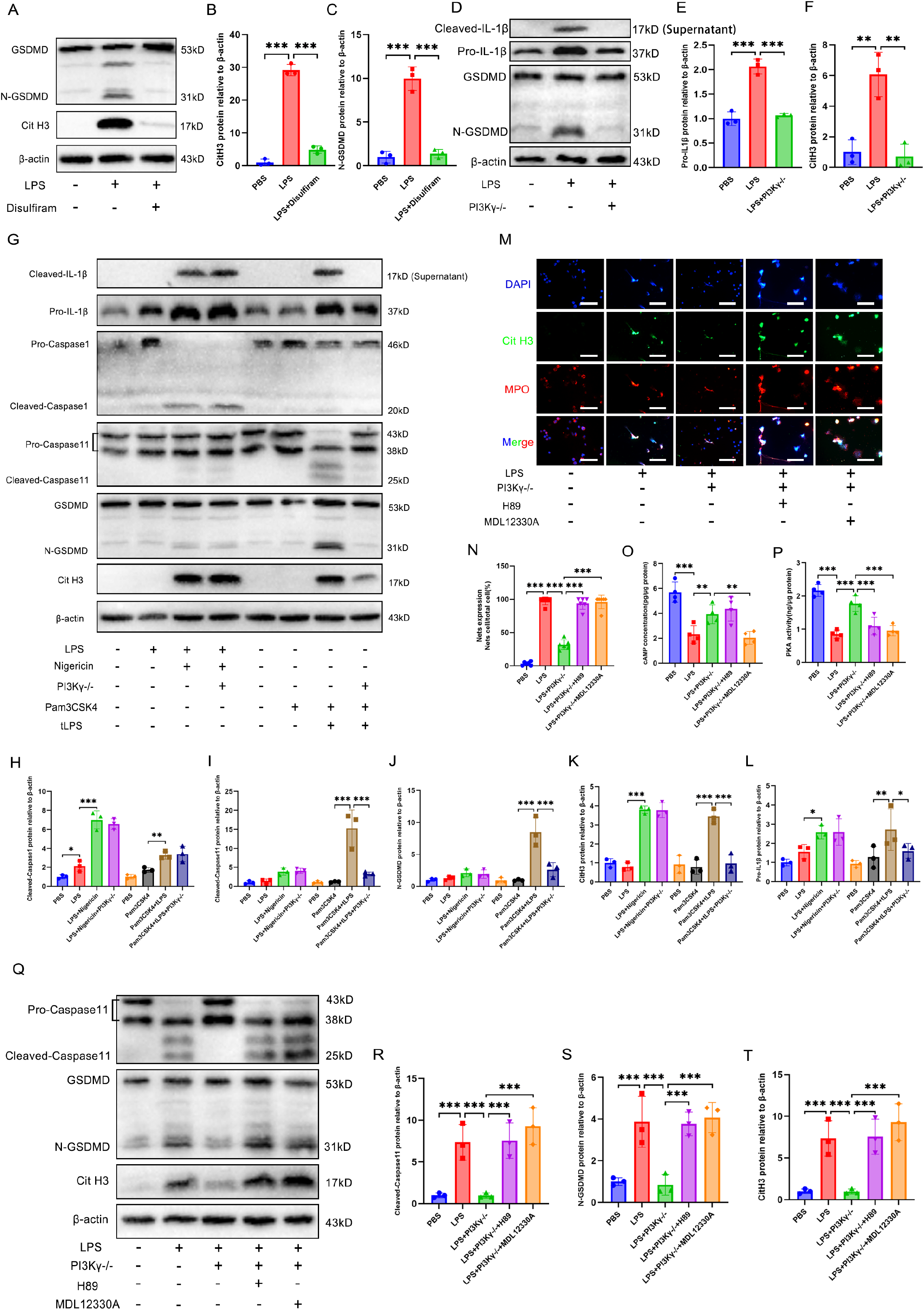
PI3Kγ promotes NETs formation via noncanonical pyroptosis pathways in vitro. (A-C) Representative western blot image (A) and quantitative comparison (B, C) of the expression of Cit H3 and GSDMD in the neutrophils of WT mice with different treatments: PBS, LPS (5μg/mL), and LPS + Disulfiram (30μM). n=3. (D-F) Representative western blot image (D) and quantitative comparison (E, F) of the expression of IL-1β and GSDMD in the neutrophils with different treatments: PBS (WT-derived neutrophils), LPS (5μg/mL, WT-derived neutrophils), LPS (PI3Kγ-/- derived neutrophil). n=3. (G-L) Representative western blot image (G) and quantitative comparison (K-L) of the expression of Cit H3, IL-1β, GSDMD, Caspase11, and Caspase1 in the neutrophils with different treatments: PBS (WT-derived neutrophils), LPS (100ng/mL, WT-derived neutrophils), LPS (100ng/mL) + Nigericin (10μM, WT-derived neutrophil), LPS (100ng/mL) + Nigericin (10μM, PI3Kγ-/- derived neutrophil); PBS (WT-derived neutrophils), Pam3CSK4 (1μg/mL, WT-derived neutrophils), Pam3CSK4 (1μg/mL) + tLPS (10μg/mL, transfer LPS, WT-derived neutrophil), Pam3CSK4 (1μg/mL) + tLPS (10μg/mL, PI3Kγ-/-derived neutrophil). n=3. (M and N) Representative immunofluorescence staining images (M) and quantitative comparison (N) of NETs produced by neutrophils with different treatments: PBS (WT-derived neutrophils), LPS (5μg/mL, WT-derived neutrophils), LPS (5μg/mL, PI3Kγ-/- derived neutrophil), LPS (5μg/mL) + H89 (20μM, PI3Kγ-/- derived neutrophil), LPS (5μg/mL) + MDL12330A (10μM, PI3Kγ-/- derived neutrophil). NETs were detected using immunofluorescent staining of DNA (DAPI, blue), Citrullinated histone 3 (Cit H3, Green), and Myeloperoxidase (MPO, red). NETs expression was calculated by NETs cell numbers/total cell numbers per high-power field (40X). n=6. Scale bars, 50 μm. (O) Comparison of cAMP concentration of neutrophil with different treatments as described in (M) by Elisa Kit. n=4. (P) Comparison of PKA kinase activity of neutrophil with different treatments as described in (M) by kinase activity assay Kit. n=4. (Q-T) Representative western blot image (Q) and quantitative comparison (R-T) of the expression of Cit H3, GSDMD, and Caspase11 in the neutrophils with different treatments as described in (M). n=3. (B-C, E-F, H-L, N-P and R-T) One-way ANOVA followed by Fisher’s LSD post hoc test.

Next, we sought to explore the molecular mechanism by which blocking PI3Kγ inhibits the nonclassical pyroptosis pathway. The most common pathway is that PI3Kγ may directly regulate PI3K/AKT signaling through its lipid kinase function which is thought to be a direct downstream of PI3Kγ. To this end, we detected the NETs expression after SC79 (a specific AKT agonists) administration. The results showed that PI3Kγ blockade reduced the AKT phosphorylation and NETs formation. Interestingly, the phosphorylation of AKT was increased but CitH3 expression was decreased after SC79 treatment (Figure S7A-S7F). These results suggested that PI3Kγ regulated noncanonical pyroptosis independent of AKT, and there may be alternative signal pathway. PI3Kγ can function as an anchor protein^35^ for protein kinase A (PKA) to regulate the cAMP/PKA signaling pathway in a negative feedback loop ^36^. We next speculated whether PKA signaling was involved in PI3Kγ-mediated regulation of noncanonical pyroptosis pathway. H89 (a PKA inhibitor) and MDL12330A (an adenylyl cyclase inhibitor, cAMP product inhibitor) were used respectively to block the PKA signaling. Immunofluorescence analysis shows that decreased NETs expression by PI3Kγ blockade was counteracted by H89 or MDL12330A (Figure 6M-6N). Moreover, cAMP/PKA signal was evaluated by ELISA and kinase active assay. The results suggested that PI3Kγ inhibition reversed the LPS-induced activation of cAMP/PKA signaling, while H89 and MDL12330A attenuated such processes (Figure 6O-6P). Concomitantly, PI3Kγ deficiency deactivated LPS-induced Caspase 11/GSDMD signaling pathway. Interestingly, PI3Kγ-mediated inhibition of noncanonical pyroptosis in neutrophils was markedly reversed by pre-treatment of H89 or MDL12330A (Figure 6Q-6T). Similarly, these results above were recapitulated in IPI549-treated neutrophils from WT mice (Figure S8A-S8F). Together, these data indicate that PI3Kγ may regulate noncanonical pyroptosis through the cAMP/PKA signaling pathway.

### cAMP/PKA inhibitor eliminates protective effect of PI3Kγ knockout in elastase-induced AAA

To investigate the possible involvement of cAMP/PKA signaling in AAA, PI3Kγ-/- mice received PPE-induced AAA and H89. As shown in Figure 7A, H89 treatment significantly increased the AAA size compared with vehicle treatment (Figure 7B-7E). The elastin fragmentation in the aortic wall was more severe in H89 group than in the vehicle group (Figure 7F-7G). Further analysis revealed that H89 significantly decreased the mRNA levels of Acta2, Cnn1, Tagln and Myh11 in the abdominal aorta and increased the mRNA levels of Klf4. The mRNA levels of IL6, TNFa, Ccl2, IL-1β and MMP2 were also upregulated after H89 administration (Figure 7H). Then, immunofluorescence and Western blot results showed a significant increase in the expression of NETs with H89 administration compared with that in vehicle group (Figure 7I-7J and 7O-7P). Moreover, H89 aggravated Caspase 11 activation and GSDMD cleavage in the arterial wall of PI3Kγ-/- mice (Figure 7I and 7K-7L). Additionally, cAMP concentration and PKA activity were also detected. H89 treatment significantly inhibited PKA kinase activity, while no obvious increase in cAMP levels was observed (Figure 7M-7N). These results suggested that PI3Kγ deficiency suppressed noncanonical pyroptosis thus inhibited NET formation in a cAMP/PKA-dependent manner.

**Figure 7.**
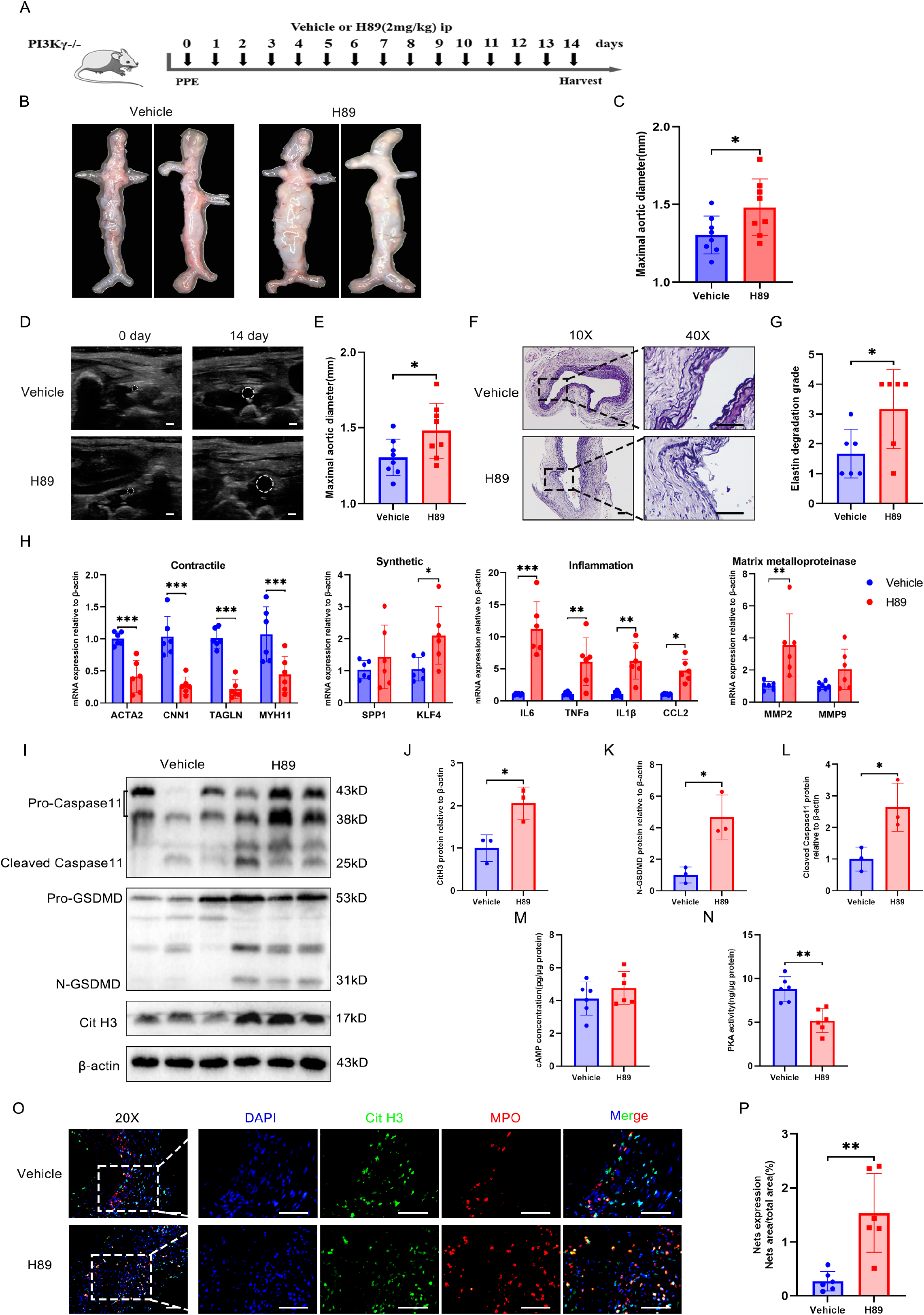
cAMP/PKA inhibitor eliminates protective effect of PI3Kγ knockout in elastase-induced AAA (A) Schematic diagram of the animal study to verify the effect of H89 administration in PPE-induced AAA PI3Kγ-/- mice. (B and C) Representative macroscopic images (B) and quantitative comparison (C) of maximal aortic diameter in PI3Kγ-/- mice after PPE surgery. n=8. (D and E) Representative ultrasonic images (D) and quantitative comparison (E) of maximal lumen diameter in PI3Kγ-/- mice after PPE surgery. n=8. Scale bar: 1 mm. (F and G) Representative VVG staining images (F) and quantitative comparison (G) of VVG staining score in PI3Kγ-/- mice after PPE surgery. n = 6. Scale bars, 50 μm. (H) Relative mRNA levels of contractile, synthetic, inflammation, and matrix metalloproteinase genes in the abdominal aorta of PI3Kγ-/- mice. n=6. (I-L) Representative western blot image (I) and quantitative analysis (J-L) of Cit H3, GSDMD, and Caspase11 protein expression in the abdominal aorta of PI3Kγ-/- mice. n = 3. (M and N) The concentration of cAMP (M) and PKA activity (N) in the abdominal aorta of PI3Kγ-/- mice. n = 6. (O and P) Representative immunofluorescence staining images (O) and quantitative comparison (P) of NETs in PI3Kγ-/- mice abdominal aorta after PPE-induced AAA. Nets expression was calculated by NET area/total area (%) according to the fluorescence colocalization of DNA (DAPI, Blue), Citrullinated histone 3 (Cit H3, Green), and Myeloperoxidase (MPO, red). n=6. Scale bars, 50 μm (C, E, G, J-N and P) Two-tailed unpaired student’s t test. (H) Two-way ANOVA followed by Bonferroni test.

## Discussion

Robust infiltration of inflammatory cells plays a critical role in the pathogenesis of AAA. Among those inflammatory responses, NETs derived by neutrophils are a key trigger for inducing AAA dilation^6^. Therefore, targeting NETs formation may represent a new avenue for treating AAA. Here, we revealed that PI3Kγ, the upstream regulatory molecule of NETs formation, may be a potential target for inhibiting AAA progression. Meanwhile, we elucidated that the molecular mechanism of PI3Kγ regulating NETs release is dependent on noncanonical pyroptosis. These findings provide a theoretical basis for targeting PI3Kγ to reduce arterial wall inflammation and treat AAA.

A previous study found that NETs induced VSMC apoptosis through p38/JNK signaling, degraded elastic fibers, and promoted Ang II-induced AAA^37^. Another recent study showed that circulating NETs were positively correlated with clinical outcomes of AAA; NETs promoted VSMC inflammatory release and phenotypic transformation through Hippo-Yap signaling activation, exacerbating AAA dilation^38^. These publications suggest that NETs play a critical role in promoting AAA progression. We also found a significant increase in NETs formation in human AAA and mouse elastase-induced AAA, which is consistent with their findings. Meanwhile, we determined the expression of NETs at different time points in mouse model of elastase-induced AAA; data showed that the NETs level reached at peak during day 3 to day 7, and mildly descended at day 14 after PPE surgery. This is consistent with previous studies which show that neutrophils are the initiating cells of the inflammatory response and infiltration of the arterial wall occurs in the early stage of aneurysm formation, which may explain the decrease in NETs levels in the late stage of aneurysm.

To date, studies on NETs formation and aneurysm remain scarce. Although several studies have revealed the causal relationship between NETs and AAA^37–39^, they mainly focused on the downstream mechanism by which NETs promoted AAA formation, while rarely discussed its upstream regulatory mechanisms which may be the key target for therapeutic drugs discovery. A recent study reported that treatment with GSK484, a PAD4 inhibitor, inhibited NETs formation and attenuated AngII-induced experimental aneurysm progression^16^. We subsequently investigated the protective effect of Cl amidine (a specific PAD4 inhibitor) on aneurysms, and the results are consistent with previous studies^20^, suggesting that NETs formation is involved deeply during AAA formation. Unfortunately, application of PAD inhibitors targeting NETs formation will damage the innate immune function^22^ and is not suitable for prolonged administration in treatment of AAA. Therefore, the identification of upstream regulators of NETs is crucial for AAA treatment. Previous studies have shown that inhibition of PI3Kγ promotes neutrophil senescence and reduces NETs, which in turn treats acute lung injury^40^. PI3Kγ depletion protects against microscopic polyangiitis by inhibiting NETs formation^26^. We hypothesized that PI3Kγ may be involved in AAA progression by affecting NETs formation. In this study, we identified that PI3Kγ was significantly upregulated in elastase-induced AAA mice, and PI3Kγ knockout significantly inhibited the formation of AAA. Furthermore, PI3Kγ blockade dramatically suppressed neutrophil infiltration and NETs formation, inhibiting AAA formation. These suggest that PI3Kγ may be the upstream regulatory molecule of NETs in AAA formation and progression.

PI3Kγ is widely expressed in many inflammatory cell types^41^. Previous studies suggested that PI3Kγ promoted carotid re-endothelialization and aggravated vascular stenosis through Cxcl10 secretion in Th1 cells^42^. In macrophages, PI3Kγ facilitates LDL uptake to form foam cells and exacerbated atherosclerotic plaque formation^43^. Given that neutrophils are the most abundant immune cell type and neutrophil infiltration is involved in AAA formation, we hypothesized that PI3Kγ may also promote the release of NETs and aggravate AAA. To clarify whether it is neutrophil-derived PI3Kγ that regulates NETs release and promotes AAA, we isolated neutrophils from mice bone marrow. Our data suggests that inhibition of PI3Kγ reduced NETs release, which is consistent with previous studies^44^. Subsequently, bone marrow neutrophils from WT mice were adoptive transferred to PI3Kγ-/- mice to precisely dissect the contribution of PI3Kγ in neutrophil in elastase-induced AAA. The results showed that adoptively transferred WT neutrophils significantly worsened AAA in PI3Kγ-/- mice, elucidating that neutrophil-derived PI3Kγ is involved in aneurysms.

Despite the fact that PI3Kγ is involved in regulating NETs formation during AAA, the molecular mechanism is still poorly understood. Previous studies have shown that reactive oxygen species (ROS) is an important inducer regulating the formation of NETs^45^, and NADPH oxidase enzyme 2 (NOX2) is the key enzyme that produces ROS to mediate the formation of NETs^46^. However, our data showed that PI3Kγ blockade reduced the expression of NETs, but not decreased the upregulation of NOX2 upon LPS stimulation, and had a synergistic effect when combined with NOX2 inhibitor. This suggested that PI3Kγ regulated NETs expression independent of ROS signaling pathways. Recent studies have shown that GSDMD, as the key executioner of pyroptosis, is required for the formation of NETs^33^. Cleaved GSDMD enhances nuclear membrane permeability at the early stage of NETs formation, thus facilitates the entry of cytoplasmic granules into the nucleus followed by pore formation at the cytoplasmic membrane to promote NETs complex release^47^. The FDA-approved drug disulfiram (DSF), which is used to treat alcohol addiction, recently shows the ability to inhibit GSDMD pore formation^48^. We found that LPS-induced NETs formation was significantly reduced after DSF administration. This is consistent with previous studies^49^. We therefore hypothesized that PI3Kγ regulated NETs expression through pyroptosis. We found that the pyroptosis-related N-GSDMD was significantly increased in human AAA, which proved the clinical relevance of pyroptosis in AAA. In addition, pyroptosis signal was significantly reduced under PI3Kγ deficiency, revealing that PI3Kγ was involved in the regulation of pyroptosis. This is the first time that we report a link between PI3Kγ and pyroptosis.

However, mechanistic insights into the effects of PI3Kγ on pyroptosis are unknown. Previous studies have shown that the induction of pyroptosis can be mediated by two pathways: canonical and noncanonical pyroptosis. Canonical pyroptosis results from activation of caspase1 by the inflammatory complex, whereas noncanonical pyroptosis results from activation of caspase11 by intracellular LPS ^34^. Thus, we induced canonical and noncanonical pyroptosis using LPS prime followed by nigericin stimulation and Pam3CSK4 prime followed by LPS transfection, respectively according to the previous publications^50^. The results showed that PI3Kγ deficiency reduced the formation of NETs, and inhibited the activation of noncanonical pyroptosis and the release of IL-1β, but had no effect on the canonical pyroptosis pathway. These data suggest that PI3Kγ regulate NETs formation via noncanonical pyroptosis.

To further explore the mechanism of PI3Kγ regulating noncanonical pyroptosis, the following attempts were made. The first thing that comes to mind is that PI3Kγ has phosphoinositide kinase activity which phosphorylates phosphatidylinositol 2 phosphate PIP2 to phosphatidylinositol 3 phosphate PIP3; phosphorylation of PIP3 recruits AKT with a PH structural domain and activates downstream signaling. Previous studies indicated that PI3Kγ drived NETs formation dependent on the activation of AKT signaling^40, 44^. However, they did not perform rescue experiments to prove that PI3Kγ regulates NETs formation through PI3K/AKT. Our results suggest that PI3Kγ blockade reduced NETs formation and was accompanied by AKT signaling inhibition, which was consistent with their results. In contrast, when AKT signaling was significantly activated by SC79, NETs levels were further reduced, implying that blocking PI3Kγ and AKT activation play a synergistic role in suppressing NETs formation. A recent study suggests that the activation of AKT/mTOR autophagy pathway promoted the transformation of neutrophil function from NETosis to phagocytosis^51^. This may explain the protective effect on NETs formation after AKT signaling activation. The above results indicated that PI3Kγ regulated NETs in a manner independent of AKT signaling.

Patrucco et al. found that PI3Kγ, in addition to phosphatidyl kinase activity, acted as an anchoring protein to exert a bridging function, negatively regulating cardiac contractility^52^. PI3Kγ promoted the interaction of PKA with PDEs and negatively regulates cAMP/PKA signaling^53^. We hypothesized that PI3Kγ regulated neutrophil noncanonical pyroptosis through cAMP/PKA signaling, The results showed that the cAMP/PKA signaling activated by PI3Kγ deletion was re-suppressed in neutrophils after treatment with H89 and MDL12330A, and the protective effect of PI3Kγ deletion on NETs formation was abolished. Additionally, the inhibition of noncanonical pyroptosis signaling by PI3Kγ deficiency was also reversed after H89 and MDL12330A administration. Furthermore, we also explored the role of cAMP/PKA signaling in PI3Kγ-mediated AAA within elastase-induced AAA. The results showed that blocking PKA signaling abrogated the protective effect of PI3Kγ deficiency on AAA, while NETs formation and noncanonical pyroptosis signaling in the arterial wall were also significantly enhanced. These results explain the molecular mechanism of PI3Kγ regulating noncanonical pyroptosis mediated NETs formation in AAA.

This study has certain limitations. Although we have showed that PI3Kγ inhibition reduces NETs formation, precise molecular mechanism of NETs formation remains unclear. Since there are different stimuli-specific signaling pathways during NETosis, further investigations should be performed on the potential crosstalk between various upstream signaling pathways of NETosis, especially in AAA field.

In conclusion, our results revealed the molecular mechanism of PI3Kγ in regulating NETs formation. Meanwhile, we found that PI3Kγ blockade significantly inhibit neutrophil infiltration and NETs formation in the abdominal aorta, notably ameliorate multiple pathological changes in AAA. Therefore, the results of this study help to promote PI3Kγ inhibitor represents a useful candidate for development of drugs against AAA.

## Author contributions

BH. P., and W.W. designed the study and revised the paper. YC. X. performed experiments and wrote the paper. S.L., JN.Z., and JJ.S. helped with animal experiments. Y.L. contributed to sample collection.

---

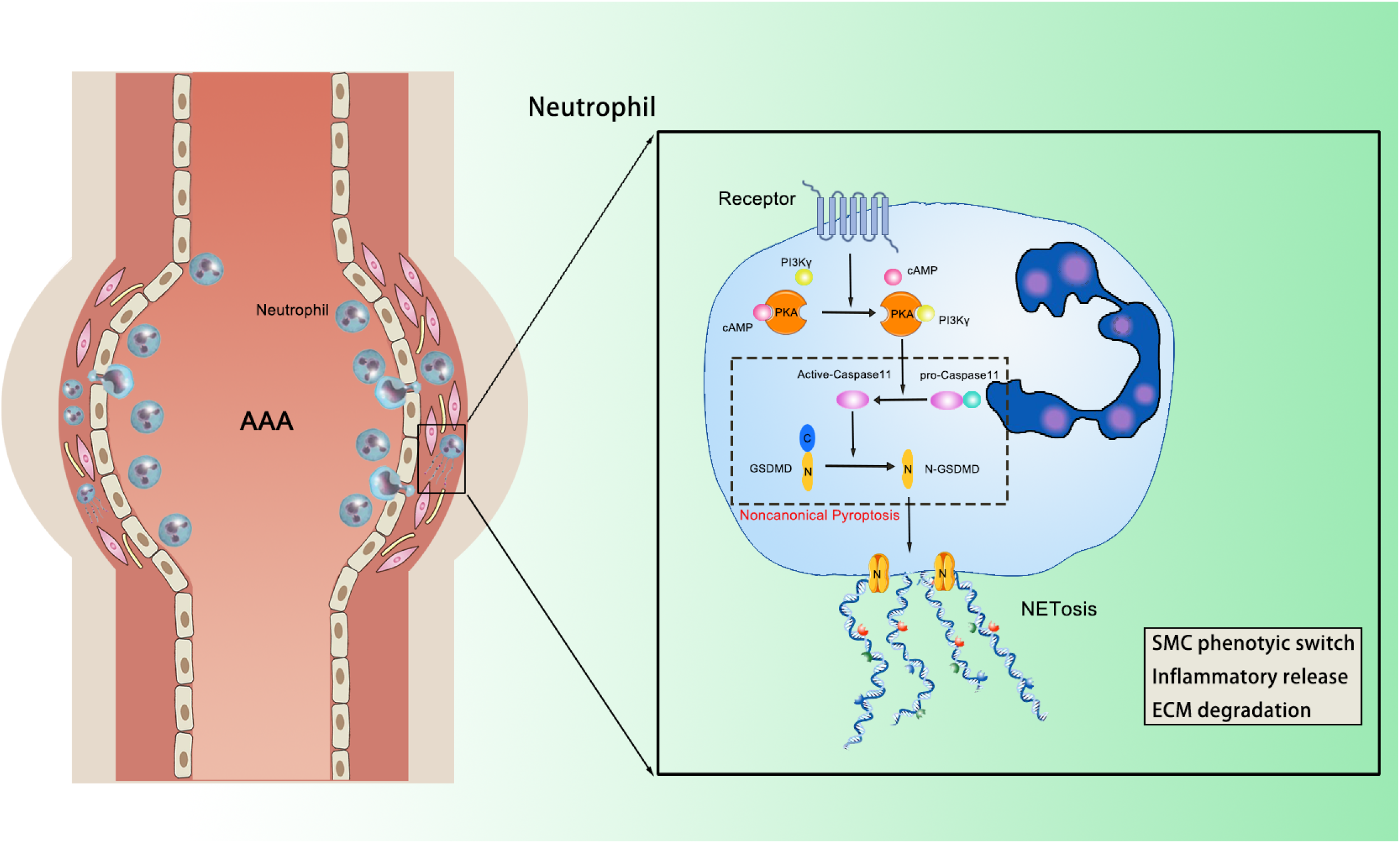

## Reference

1. Baman JR, Eskandari MK. What is an abdominal aortic aneurysm? Jama. 2022;328:2280

2. Song P, He Y, Adeloye D, Zhu Y, Ye X, Yi Q, Rahimi K, Rudan I. The global and regional prevalence of abdominal aortic aneurysms: A systematic review and modelling analysis. Ann Surg. 2022

3. Guirguis-Blake JM, Beil TL, Senger CA, Coppola EL. Primary care screening for abdominal aortic aneurysm: Updated evidence report and systematic review for the us preventive services task force. Jama. 2019;322:2219–2238

4. Golledge J, Moxon JV, Singh TP, Bown MJ, Mani K, Wanhainen A. Lack of an effective drug therapy for abdominal aortic aneurysm. J Intern Med. 2020;288:6–22

5. Wang W, Xu B, Xuan H, Ge Y, Wang Y, Wang L, Huang J, Fu W, Michie SA, Dalman RL. Hypoxia-inducible factor 1 in clinical and experimental aortic aneurysm disease. J Vasc Surg. 2018;68:1538–1550.e1532

6. Yuan Z, Lu Y, Wei J, Wu J, Yang J, Cai Z. Abdominal aortic aneurysm: Roles of inflammatory cells. Front Immunol. 2020;11:609161

7. Quintana RA, Taylor WR. Cellular mechanisms of aortic aneurysm formation. Circ Res. 2019;124:607–618

8. Eliason JL, Hannawa KK, Ailawadi G, Sinha I, Ford JW, Deogracias MP, Roelofs KJ, Woodrum DT, Ennis TL, Henke PK, Stanley JC, Thompson RW, Upchurch GR, Jr. Neutrophil depletion inhibits experimental abdominal aortic aneurysm formation. Circulation. 2005;112:232–240

9. Parikh RR, Folsom AR, Poudel K, Lutsey PL, Demmer RT, Pankow JS, Chen LY, Tang W. Association of differential leukocyte count with incident abdominal aortic aneurysm over 22.5 years: The aric study. Arterioscler Thromb Vasc Biol. 2021;41:2342–2351

10. Pagano MB, Bartoli MA, Ennis TL, Mao D, Simmons PM, Thompson RW, Pham CT. Critical role of dipeptidyl peptidase i in neutrophil recruitment during the development of experimental abdominal aortic aneurysms. Proc Natl Acad Sci U S A. 2007;104:2855–2860

11. Klopf J, Brostjan C, Neumayer C, Eilenberg W. Neutrophils as regulators and biomarkers of cardiovascular inflammation in the context of abdominal aortic aneurysms. Biomedicines. 2021;9

12. Boivin G, Faget J, Ancey PB, Gkasti A, Mussard J, Engblom C, Pfirschke C, Contat C, Pascual J, Vazquez J, Bendriss-Vermare N, Caux C, Vozenin MC, Pittet MJ, Gunzer M, Meylan E. Durable and controlled depletion of neutrophils in mice. Nat Commun. 2020;11:2762

13. Poli V, Zanoni I. Neutrophil intrinsic and extrinsic regulation of netosis in health and disease. Trends Microbiol. 2022

14. Herre M, Cedervall J, Mackman N, Olsson AK. Neutrophil extracellular traps in the pathology of cancer and other inflammatory diseases. Physiol Rev. 2023;103:277–312

15. Remijsen Q, Kuijpers TW, Wirawan E, Lippens S, Vandenabeele P, Vanden Berghe T. Dying for a cause: Netosis, mechanisms behind an antimicrobial cell death modality. Cell Death Differ. 2011;18:581–588

16. Eilenberg W, Zagrapan B, Bleichert S, Ibrahim N, Knöbl V, Brandau A, Martelanz L, Grasl MT, Hayden H, Nawrozi P, Rajic R, Häusler C, Potolidis A, Schirwani N, Scheuba A, Klopf J, Teubenbacher P, Weigl MP, Kirchweger P, Beitzke D, Stiglbauer-Tscholakoff A, Panzenböck A, Lang I, Mauracher LM, Hell L, Pabinger I, Bailey MA, Scott DJA, Maegdefessel L, Busch A, Huk I, Neumayer C, Brostjan C. Histone citrullination as a novel biomarker and target to inhibit progression of abdominal aortic aneurysms. Transl Res. 2021;233:32–46

17. Yang S, Chen L, Wang Z, Chen J, Ni Q, Guo X, Liu W, Lv L, Xue G. Neutrophil extracellular traps induce abdominal aortic aneurysm formation by promoting the synthetic and proinflammatory smooth muscle cell phenotype via hippo-yap pathway. Transl Res. 2022

18. Plana E, Oto J, Medina P, Fernández-Pardo Á, Miralles M. Novel contributions of neutrophils in the pathogenesis of abdominal aortic aneurysm, the role of neutrophil extracellular traps: A systematic review. Thromb Res. 2020;194:200–208

19. Spinosa M, Su G, Salmon MD, Lu G, Cullen JM, Fashandi AZ, Hawkins RB, Montgomery W, Meher AK, Conte MS, Sharma AK, Ailawadi G, Upchurch GR, Jr. Resolvin d1 decreases abdominal aortic aneurysm formation by inhibiting netosis in a mouse model. J Vasc Surg. 2018;68:93s–103s

20. Meher AK, Spinosa M, Davis JP, Pope N, Laubach VE, Su G, Serbulea V, Leitinger N, Ailawadi G, Upchurch GR, Jr. Novel role of il (interleukin)-1β in neutrophil extracellular trap formation and abdominal aortic aneurysms. Arterioscler Thromb Vasc Biol. 2018;38:843–853

21. Liu X, Arfman T, Wichapong K, Reutelingsperger CPM, Voorberg J, Nicolaes GAF. Pad4 takes charge during neutrophil activation: Impact of pad4 mediated net formation on immune-mediated disease. J Thromb Haemost. 2021;19:1607–1617

22. Li P, Li M, Lindberg MR, Kennett MJ, Xiong N, Wang Y. Pad4 is essential for antibacterial innate immunity mediated by neutrophil extracellular traps. J Exp Med. 2010;207:1853–1862

23. Márquez-Sánchez AC, Koltsova EK. Immune and inflammatory mechanisms of abdominal aortic aneurysm. Front Immunol. 2022;13:989933

24. Zhu J, Li K, Yu L, Chen Y, Cai Y, Jin J, Hou T. Targeting phosphatidylinositol 3-kinase gamma (pi3kγ): Discovery and development of its selective inhibitors. Med Res Rev. 2021;41:1599–1621

25. DeSouza-Vieira T, Guimarães-Costa A, Rochael NC, Lira MN, Nascimento MT, Lima-Gomez PS, Mariante RM, Persechini PM, Saraiva EM. Neutrophil extracellular traps release induced by leishmania: Role of pi3kγ, erk, pi3kσ, pkc, and [ca2+]. J Leukoc Biol. 2016;100:801–810

26. Kimura H, Matsuyama Y, Araki S, Koizumi A, Kariya Y, Takasuga S, Eguchi S, Nakanishi H, Sasaki J, Sasaki T. The effect and possible clinical efficacy of in vivo inhibition of neutrophil extracellular traps by blockade of pi3k-gamma on the pathogenesis of microscopic polyangiitis. Mod Rheumatol. 2018;28:530–541

27. Chaikof EL, Dalman RL, Eskandari MK, Jackson BM, Lee WA, Mansour MA, Mastracci TM, Mell M, Murad MH, Nguyen LL, Oderich GS, Patel MS, Schermerhorn ML, Starnes BW. The society for vascular surgery practice guidelines on the care of patients with an abdominal aortic aneurysm. J Vasc Surg. 2018;67:2–77.e72

28. Liu S, Huang T, Liu R, Cai H, Pan B, Liao M, Yang P, Wang L, Huang J, Ge Y, Xu B, Wang W. Spermidine suppresses development of experimental abdominal aortic aneurysms. J Am Heart Assoc. 2020;9:e014757

29. Jakobs C, Bartok E, Kubarenko A, Bauernfeind F, Hornung V. Immunoblotting for active caspase-1. Methods Mol Biol. 2013;1040:103–115

30. Welch HC, Condliffe AM, Milne LJ, Ferguson GJ, Hill K, Webb LM, Okkenhaug K, Coadwell WJ, Andrews SR, Thelen M, Jones GE, Hawkins PT, Stephens LR. P-rex1 regulates neutrophil function. Curr Biol. 2005;15:1867–1873

31. Tong Y, Xin Y, Fu L, Shi J, Sun Y. Excessive neutrophil extracellular trap formation induced by porphyromonas gingivalis lipopolysaccharide exacerbates inflammatory responses in high glucose microenvironment. Front Cell Infect Microbiol. 2023;13:1108228

32. Chen F, Chu C, Wang X, Yang C, Deng Y, Duan Z, Wang K, Liu B, Ji W, Ding W. Hesperetin attenuates sepsis-induced intestinal barrier injury by regulating neutrophil extracellular trap formation via the ros/autophagy signaling pathway. Food Funct. 2023;14:4213–4227

33. Sollberger G, Choidas A, Burn GL, Habenberger P, Di Lucrezia R, Kordes S, Menninger S, Eickhoff J, Nussbaumer P, Klebl B, Krüger R, Herzig A, Zychlinsky A. Gasdermin d plays a vital role in the generation of neutrophil extracellular traps. Sci Immunol. 2018;3

34. Burdette BE, Esparza AN, Zhu H, Wang S. Gasdermin d in pyroptosis. Acta Pharm Sin B. 2021;11:2768–2782

35. Lanahan SM, Wymann MP, Lucas CL. The role of pi3kγ in the immune system: New insights and translational implications. Nat Rev Immunol. 2022;22:687–700

36. Perino A, Ghigo A, Ferrero E, Morello F, Santulli G, Baillie GS, Damilano F, Dunlop AJ, Pawson C, Walser R, Levi R, Altruda F, Silengo L, Langeberg LK, Neubauer G, Heymans S, Lembo G, Wymann MP, Wetzker R, Houslay MD, Iaccarino G, Scott JD, Hirsch E. Integrating cardiac pip3 and camp signaling through a pka anchoring function of p110γ. Mol Cell. 2011;42:84–95

37. Wei M, Wang X, Song Y, Zhu D, Qi D, Jiao S, Xie G, Liu Y, Yu B, Du J, Wang Y, Qu A. Inhibition of peptidyl arginine deiminase 4-dependent neutrophil extracellular trap formation reduces angiotensin ii-induced abdominal aortic aneurysm rupture in mice. Front Cardiovasc Med. 2021;8:676612

38. Yang S, Chen L, Wang Z, Chen J, Ni Q, Guo X, Liu W, Lv L, Xue G. Neutrophil extracellular traps induce abdominal aortic aneurysm formation by promoting the synthetic and proinflammatory smooth muscle cell phenotype via hippo-yap pathway. Transl Res. 2023;255:85–96

39. Chen L, Liu Y, Wang Z, Zhang L, Xu Y, Li Y, Zhang L, Wang G, Yang S, Xue G. Mesenchymal stem cell-derived extracellular vesicles protect against abdominal aortic aneurysm formation by inhibiting net-induced ferroptosis. Exp Mol Med. 2023;55:939–951

40. Song D, Adrover JM, Felice C, Christensen LN, He XY, Merrill JR, Wilkinson JE, Janowitz T, Lyons SK, Egeblad M, Tonks NK. Ptp1b inhibitors protect against acute lung injury and regulate cxcr4 signaling in neutrophils. JCI Insight. 2022;7

41. Kaneda MM, Messer KS, Ralainirina N, Li H, Leem CJ, Gorjestani S, Woo G, Nguyen AV, Figueiredo CC, Foubert P, Schmid MC, Pink M, Winkler DG, Rausch M, Palombella VJ, Kutok J, McGovern K, Frazer KA, Wu X, Karin M, Sasik R, Cohen EE, Varner JA. Pi3kγ is a molecular switch that controls immune suppression. Nature. 2016;539:437–442

42. Lupieri A, Smirnova NF, Solinhac R, Malet N, Benamar M, Saoudi A, Santos-Zas I, Zeboudj L, Ait-Oufella H, Hirsch E, Ohayon P, Lhermusier T, Carrié D, Arnal JF, Ramel D, Gayral S, Laffargue M. Smooth muscle cells-derived cxcl10 prevents endothelial healing through pi3kγ-dependent t cells response. Cardiovasc Res. 2020;116:438–449

43. Anzinger JJ, Chang J, Xu Q, Barthwal MK, Bohnacker T, Wymann MP, Kruth HS. Murine bone marrow-derived macrophages differentiated with gm-csf become foam cells by pi3kγ-dependent fluid-phase pinocytosis of native ldl. J Lipid Res. 2012;53:34–42

44. de Carvalho Oliveira V, Tatsiy O, McDonald PP. Phosphoinositol 3-kinase-driven net formation involves different isoforms and signaling partners depending on the stimulus. Front Immunol. 2023;14:1042686

45. Zhan X, Wu R, Kong XH, You Y, He K, Sun XY, Huang Y, Chen WX, Duan L. Elevated neutrophil extracellular traps by hbv-mediated s100a9-tlr4/rage-ros cascade facilitate the growth and metastasis of hepatocellular carcinoma. Cancer Commun (Lond). 2023;43:225–245

46. Lv X, Liu Z, Xu L, Song E, Song Y. Tetrachlorobenzoquinone exhibits immunotoxicity by inducing neutrophil extracellular traps through a mechanism involving ros-jnk-nox2 positive feedback loop. Environ Pollut. 2021;268:115921

47. Chen KW, Monteleone M, Boucher D, Sollberger G, Ramnath D, Condon ND, von Pein JB, Broz P, Sweet MJ, Schroder K. Noncanonical inflammasome signaling elicits gasdermin d-dependent neutrophil extracellular traps. Sci Immunol. 2018;3

48. Hu JJ, Liu X, Xia S, Zhang Z, Zhang Y, Zhao J, Ruan J, Luo X, Lou X, Bai Y, Wang J, Hollingsworth LR, Magupalli VG, Zhao L, Luo HR, Kim J, Lieberman J, Wu H. Fda-approved disulfiram inhibits pyroptosis by blocking gasdermin d pore formation. Nat Immunol. 2020;21:736–745

49. Silva CMS, Wanderley CWS, Veras FP, Sonego F, Nascimento DC, Gonçalves AV, Martins TV, Cólon DF, Borges VF, Brauer VS, Damasceno LEA, Silva KP, Toller-Kawahisa JE, Batah SS, Souza ALJ, Monteiro VS, Oliveira AER, Donate PB, Zoppi D, Borges MC, Almeida F, Nakaya HI, Fabro AT, Cunha TM, Alves-Filho JC, Zamboni DS, Cunha FQ. Gasdermin d inhibition prevents multiple organ dysfunction during sepsis by blocking net formation. Blood. 2021;138:2702–2713

50. Kayagaki N, Wong MT, Stowe IB, Ramani SR, Gonzalez LC, Akashi-Takamura S, Miyake K, Zhang J, Lee WP, Muszyński A, Forsberg LS, Carlson RW, Dixit VM. Noncanonical inflammasome activation by intracellular lps independent of tlr4. Science. 2013;341:1246–1249

51. Guo G, Liu Z, Yu J, You Y, Li M, Wang B, Tang J, Han P, Wu J, Shen H. Neutrophil function conversion driven by immune switchpoint regulator against diabetes-related biofilm infections. Adv Mater. 2023:e2310320

52. Patrucco E, Notte A, Barberis L, Selvetella G, Maffei A, Brancaccio M, Marengo S, Russo G, Azzolino O, Rybalkin SD, Silengo L, Altruda F, Wetzker R, Wymann MP, Lembo G, Hirsch E. Pi3kgamma modulates the cardiac response to chronic pressure overload by distinct kinase-dependent and - independent effects. Cell. 2004;118:375–387

53. Ghigo A, Perino A, Mehel H, Zahradníková A, Jr., Morello F, Leroy J, Nikolaev VO, Damilano F, Cimino J, De Luca E, Richter W, Westenbroek R, Catterall WA, Zhang J, Yan C, Conti M, Gomez AM, Vandecasteele G, Hirsch E, Fischmeister R. Phosphoinositide 3-kinase γ protects against catecholamine-induced ventricular arrhythmia through protein kinase a-mediated regulation of distinct phosphodiesterases. Circulation. 2012;126:2073–2083

